# Nuclear trafficking of *Anelloviridae* capsid protein ORF1 reflects modular evolution of subcellular targeting signals

**DOI:** 10.1101/2025.05.14.653747

**Authors:** Gayle F. Petersen, Silvia Pavan, Daryl Ariawan, Ole Tietz, Sepehr Nematollahzadeh, Subir Sarker, Jade K. Forwood, Gualtiero Alvisi

**Affiliations:** Gulbali Institute, Charles Sturt University, Wagga Wagga, NSW 2678, Australia; Department of Molecular Medicine, University of Padova, Padova 35121, Italy; Dementia Research Centre, Macquarie Medical School, Faculty of Medicine, Health and Human Sciences, Macquarie University, Sydney, NSW 2109, Australia; Biomedical Sciences & Molecular Biology, College of Medicine and Dentistry, James Cook University, Townsville, QLD 4811, Australia; School of Agricultural, Environmental and Veterinary Sciences, Charles Sturt University, Wagga Wagga, NSW 2678, Australia

**Keywords:** *Anelloviridae*, karyopherins, shuttling, nucleolus, capsid, NLS, NoLS, projection domain

## Abstract

*Anelloviridae* members are ubiquitous viruses with a small, negative sense, single-stranded DNA genome which is replicated by host cell DNA polymerases. Anelloviruses are postulated to interact with the host cell nuclear transport machinery, however, the lack of reliable cell culture models strongly limits our knowledge regarding *Anelloviridae*-host interactions. In particular, capsid nuclear import is a largely uncharacterized process. We addressed this by investigating the relationship between host cell nuclear transport receptors (NTRs) and ORF1, the putative capsid protein from torque teno douroucouli virus (TTDoV). We identified the subcellular targeting signals and NTRs responsible for its nucleolar and nuclear localization, and characterized their relative contribution to ORF1 subcellular localization. In the absence of other viral proteins, ORF1 accumulated in the nucleoli. Bioinformatics analysis revealed a putative nuclear localization signal (NLS) within the highly conserved N-terminal arginine rich motif (ARM) (“NLSn”, 27-RRWRRRPRRRRRPYR-RRPYRRYGRRRKVRRR-57), and an additional C-terminal NLS (“NLSc”, 632-LPPPEKRARWGF-643), which has been specifically acquired by *Anelloviridae* capsids with larger projection domains. Such NLSs play distinct roles in ORF1 subcellular localization. NLSn features broad importin (IMP) binding affinity yet plays a minor role in nuclear import, being responsible for nucleolar targeting likely through interaction with nucleolar components. NLSc specifically interacts with IMPα and is the main driver of active nuclear transport in an IMPα/β1-dependent fashion. These findings suggest an evolutionary correlation between the acquisition of progressively larger projection domains and the presence of additional NLSs in *Anelloviridae* capsids, aimed at maximizing IMPα/β1-mediated nuclear import.

## 1. Introduction

*Anelloviridae* are small, non-enveloped viruses with circular, single-stranded negative sense DNA genomes, first discovered in 1997 when torque teno virus (TTV) was identified in the blood of a post-transfusion patient with elevated liver enzymes using molecular techniques (1). Subsequent advances in metagenomics have led to the identification of a plethora of additional TTVs infecting humans and animals, with high genomic heterogeneity (1-3). These viruses are now classified under the *Anelloviridae* family, which comprises 31 established genera with genomes between 1.6 and 3.9 kb in length (4). Larger genomes, such as those from the genera *Alphatorquevirus* and *Zetatorquevirus* (e.g., TTVs) ranging from 3.6 to 3.9 kb, usually encode for larger capsid proteins compared to smaller genomes, such as those from the genus *Gyrovirus* ranging from 1.8 to 2.4 kb (5). All these isolates feature a shared genetic organization, with a relatively conserved untranslated region (UTR) containing a GC-rich zone, and a coding region containing the major open reading frames (ORFs), ORF1, which is proposed to encode for the capsid protein (5), and an overlapping ORF2, which is believed to be a regulatory protein that interferes with host antiviral defenses by suppressing the NF-ϰB pathway (6). Depending on the isolate, several additional ORFs have also been described (7).

TTVs are believed to be the most abundant eukaryotic viruses in the human virome (7, 8), chronically infecting most of the human population, and have been detected in up to 90% of tested individuals in the absence of clinical symptoms (9). Though no disease has been unequivocally linked to TTV infection, viremia has been shown to reflect the degree of immunosuppression and is being clinically investigated as a marker of solid organ transplant rejection (10). Asymptomatic anellovirus infections are also common in other mammals (11), as well as in chickens and several other avian species, suggesting long term virus-host co-evolution. Given their ability to establish lifelong infection, it is proposed that TTVs have evolved sophisticated mechanisms to manipulate host cell functions and immune defenses (12). Recently, the immune evasion properties of *Anelloviridae* members have been linked to the evolutionary acquisition of a highly variable projection domain between β-strands H and I of the jelly-roll fold within the capsid protein, which appear considerably variable in length and larger than their *Circoviridae* counterparts (5, 13). Consistent with this hypothesis, cryogenic electron microscopy (cryo-EM) analysis of virus-like particles (VLPs) of ORF1 from *Betatorquevirus* isolate LY1 revealed that the variable projection domain protrudes outside the capsid structure at the 5-fold symmetry axis, potentially exposed for recognition by neutralizing antibodies (14).

With a circular ssDNA genome of approximately 3000 bases, TTV DNA replication has long been thought to occur through a rolling circle mechanism in the nucleus of infected cells (15, 16). Therefore, TTV proteins are expected to interact with host nuclear transport receptors (NTRs), specifically importin (IMP) superfamily members, which recognize nuclear localization signals (NLSs) on cargoes to mediate their nuclear import (17, 18). While molecules smaller than ~70 kDa can passively diffuse through the nuclear pore complex (18), larger ones and those that need to quickly accumulate in the cell nucleus require energy dependent transport (19). Further, several proteins from both DNA and RNA viruses need to be actively translocated into the nucleus to foster viral replication and manipulate cell function (20, 21). TTV capsid proteins feature a highly basic arginine rich motif (ARM) at the N-terminus, which is conserved in capsids from closely related *Circoviridae* members and other highly divergent icosahedral viruses (22). Stretches of basic amino acids within ARMs can bind to NTRs and have been proposed to act as NLSs (23), however they are generally located inside the viral capsid (13, 14, 24), packaging viral genomes of both RNA (24) and DNA (13) viruses by electrostatic interaction (22). Analysis of TTV–host interactions have been so far limited by a lack of suitable cellular systems allowing viral replication (25). Heterogenous subcellular localization has been reported for ORF1 from different TTV species, but the NTRs responsible for nuclear targeting have not yet been identified. ORF1 from TTV genotype 6 HEL32 is mainly restricted to the cytoplasm when expressed as HA or GFP tagged fusions in the absence of any other viral protein (26), while in the case of genogroup 1 P/1C1 and suid TTV isolates TTSuV1 type 1C and TTSuV2 subtype 2A, ORF1 was primarily detected in the nucleoli (27, 28).

We set out to investigate the interaction between ORF1 proteins and the host cell nuclear transport machinery using a prototype TTV, specifically the *Zetatorquevirus* torque teno douroucouli virus (TTDoV) (16). Our study identified specific sequences that directly interact with several NTRs and are responsible for TTDoV ORF1 subcellular localization. ORF1 contains two putative NLSs. The N-terminal NLS (NLSn) largely overlaps with the ARM; it is crucial for nucleolar accumulation but poorly contributes to nuclear transport, despite its ability to bind a wide range of IMPα and IMPβ NTRs. An additional NLS at the C-terminus (NLSc) selectively binds IMPα’s and is the main driver of IMPα/β1-mediated nuclear localization. Intriguingly, while NLSn is widely conserved across *Anelloviridae* and *Circoviridae* members, NLSc is exclusively found within larger capsids from anelloviruses, correlating with large projection domains and establishing an evolutionary link between acquisition of large projection domains and presence of additional IMPα/β1-dependent NLSs at the C-terminus of capsid proteins.

## 2. Materials and Methods

### 2.1. Bioinformatics

The genomic sequence of TTDoV (isolate At-TTV3) was retrieved from GenBank (accession number: 11862897). The sequence of viral encoded ORF1 was retrieved from UniProt with the code Q9DUB7. The sequences of *Circoviridae* (29) and *Anelloviridae* (5) ORF1 proteins were retrieved from UniProt. Protein sequences were analyzed with cNLS Mapper (30) to identify putative NLSs. Structural models were predicted using the AlphaFold3 Server (31) and visualized using the PyMOL Molecular Graphics System (version 3.1.3.1; Schrödinger, LLC). Protein model interactions were analyzed using the PDBePISA (https://www.ebi.ac.uk/pdbe/pisa/) and PDBsum (https://www.ebi.ac.uk/thornton-srv/databases/pdbsum/) web tools. Phylogenetic analysis was performed using Clustal Omega (32) and standard settings.

### 2.2. Plasmids

Plasmids encoding NTRs used in binding assays include human importin alpha 1 (hIMPα1ΔIBB), mouse importin alpha 2 (mIMPα2ΔIBB), human importin alpha 3 (hIMPα3ΔIBB), human importin alpha 5 (hIMPα5ΔIBB), and human importin alpha 7 (hIMPα7ΔIBB), all truncated to remove the autoinhibitory importin beta binding (IBB) domain, and human importin beta 1 (hIMPβ1), human importin beta 2 (hIMPβ2), and human importin beta 3 (hIMPβ3). Genes were cloned into pET-30a(+) or pMCSG21 vectors. All plasmids contain an N-terminal 6x histidine tag and a tobacco etch virus (TEV) protease cleavage site, except mIMPα2ΔIBB (no TEV site). Plasmids pcDNA3.1/NT-GFP and pcDNA3.1/NT-GFP-SV40 LTA, mediating the expression of cycle 3 GFP or cycle 3 GFP fused to SV40 LTA NLS (PKKKRKV), were described previously (33). Mammalian expression plasmids encoding TTDoV ORF1 NLSs fused to the C-terminus of cycle 3 GFP were generated by annealing appropriate oligonucleotide pairs in vector pcDNA3.1/NT-GFP-TOPO® (Thermo Fisher Scientific, Monza, Italy). Plasmid mCherry-Bimax2, encoding a competitive inhibitor of the IMPα/β1 nuclear import pathway (34), was kindly gifted from Yoshihiro Yoneda and Masahiro Oka (Osaka, Japan), while plasmids pDsRed-C1-fibrillarin and pDsRed-C1-nucleolin (35) were kindly provided by Denis Archambault (University of Québec, Canada). Plasmid pEGFP-C1-ORF1, with TTDoV ORF1 placed downstream of the eGFP ORF and flanked by Gateway attB recombination sites, was synthesized (BioFab Research, Rome, Italy) and substitution derivatives were generated using the Quikchange mutagenesis kit (Agilent Technologies, Cernusco sul Naviglio (MI), Italy) with appropriate oligonucleotide pairs, as previously described (36). A list of all plasmids used in this study is available in Supplementary Table S1.

### 2.3. Peptides

N-terminal fluorescein isothiocyanate (FITC) tagged synthetic peptides ORF1 NLSn and ORF1 NLSc were synthesized as described previously (33, 37), using standard Fluorenyl methoxycarbonyl (Fmoc)-solid-phase peptide synthesis on low swell 100 – 200 mesh Wang resin (0.05 mmol reaction scale, 0.5 mmol g_-1_ loading) on a CEM Liberty Blue™ Peptide Synthesizer (CEM, USA). *Initial amino acid loading*: Wang resin (100-200 mesh; 0.65 mmol g_-1_, 77 mg, 0.05 mmol) was weighed into a 10 mL polypropylene syringe equipped with a porous polypropylene frit, which was used as the reaction vessel. The resin was washed with dichloromethane (3 × 5 mL) before being allowed to swell in dichloromethane (5 mL) for at least 0.5 h prior to loading of the first amino acid. A solution of Fmoc-AA-OH (4 equiv.) was dissolved in a mixture of dry dichloromethane (2 mL), N,N-dimethylformamide (2 mL), hydroxybenzotriazole (HOBt) (4 equiv.), and N,N’-diisopropylcarbodiimide (DIC) (4 equiv.), taken up into the syringe with resin, and stirred overnight using an orbital shaker. The resin was then capped with acetic anhydride (0.1 mL) and N,N-diisopropylethylamine (DIPEA) (0.1 mL) in dichloromethane (3 mL) for 30 min. The resin was then washed with dichloromethane (3 × 4 mL) and N,N-dimethylformamide (DMF) (3 × 4 mL). *Automated peptide synthesizer:* Resin was pre-swelled in 50/50 DMF and dichloromethane (DCM) for 1 hr. Amino acids were dissolved in DMF at a concentration of 0.2 M before being transferred to the synthesizer. Peptides were synthesized using sequential amid coupling from C- to N-terminus for 5 min at 90°C, using five equivalents of amino acid with 10 equivalents of activator (0.5 M DIC (N,N′-Diisopropylcar-bodiimide) in DMF) and 5 equivalents of activator base (0.5 M Oxyma (Ethyl cyanohydroxyiminoacetate), 0.05 M DIPEA (N,N-Diisopropylethylamine) in DMF), followed by Fmoc deprotection in 20% piperidine in DMF for 3 min at 75°C and 3x resin wash in DMF. Following final Fmoc deprotection, resin was removed from the synthesizer, transferred to a syringe fitted with a propylene filter, and labelled with FITC (*see below*). Double couplings were performed for arginine residues to ensure complete coupling. *FITC labelling:* Fmoc-6-aminohexanoic acid (Ahx) was coupled to the N-terminus of peptides and deprotected using standard amino acid coupling conditions. Peptide bound resin was removed from the synthesizer and transferred to a syringe fitted with a propylene filter. FITC (2 equiv.), HOBt (3 equiv.), benzotriazol-1-yloxytripyrrolidinophosphonium hexafluorophosphate (PyBOP) (3 equiv.), and DIPEA (6 equiv.) in DMF (4 mL) was taken up in the syringe and agitated overnight in an orbital shaker. Resin was then washed with DMF (x3), DCM (x3), and methanol (x3) and proceeded to cleavage. *Cleavage*: Peptide was cleaved from the resin using a cleavage cocktail of 92.5% TFA (trifluoroacetic acid), 2.5% TIPS (triisopropylsilane), 2.5% thioanisole, and 2.5% H_2_O for at least 3 hrs at room temperature, precipitated in ice cold diethyl ether, dissolved in H_2_O, and freeze dried. Peptide ORF1 NLSc was purified using a Shimadzu LC-20AD high-performance liquid chromatography (HPLC, Shimadzu, Japan). Mass spectra were obtained on a Shimadzu LCMS-8050 LCMS system (Shimadzu, Japan) in positive electron spray [ESI+] mode, fitted with a Polaris 3 C18-A 50 x 4.6 mm column (Agilent Technologies, USA). Peptide ORF1 NLSn could not be purified due to the high arginine content. FITC tagged synthetic peptides ORF1 NLSn_A and ORF1 NLSn_B were synthesized by GenScript. A list of all peptides used in this study is available in Supplementary Table S2.

### 2.4. Expression and purification of recombinant proteins

Plasmids encoding NTRs were transformed into BL21(DE3)pLysS *E. coli* cells and proteins were expressed for 24-30 hrs at 25°C using the auto-induction method (38). Bacterial cells were pelleted via centrifugation at ~7,500 *x* g at 4°C and resuspended in His buffer (50 mM phosphate buffer, 300 mM sodium chloride, 20 mM imidazole, pH 8). Cells were lysed by three freeze-thaw cycles, followed by incubation with 20 mg/mL lysozyme (Thermo Fisher Scientific, Waltham, MA, USA) and 50 mg/mL DNase (Sigma-Aldrich, St. Louis, MO, USA). Soluble extract was isolated by centrifugation at 30,000 *x* g at 4°C, clarified by 0.45 µm filtration, and injected onto a pre-equilibrated 5 mL HisTrap HP column (Cytiva, Marlborough, MA, USA). The column was washed with 20 column volumes of His buffer, and protein was eluted using a linear gradient of 20-500 mM imidazole. Peak fractions were pooled and the 6x histidine tag was cleaved by incubation with TEV protease at 4°C overnight (except mIMPα2ΔIBB; no TEV site). Proteins were further purified by size exclusion chromatography (SEC) using a pre-equilibrated HiLoad 26/600 Superdex 75 pg or HiLoad 26/600 Superdex 200 pg column (Cytiva, Marlborough, MA, USA) and SEC buffer (50 mM Tris base, 125 mM sodium chloride, pH 8). Peak fractions were pooled and run through a 5 mL HisTrap HP column, pre-equilibrated with SEC buffer, to remove uncleaved target protein and TEV protease. Flowthrough fractions were pooled and concentrated using an Amicon 10 kDa MWCO ultra centrifugal filter (Merck Millipore, Burlington, MA, USA). Purified NTR protein was flash frozen in liquid nitrogen, aliquoted, and stored at −80°C for future use.

### 2.5. Electrophoretic mobility shift assays (EMSAs)

To qualitatively assess the interaction between NLSs and NTRs, EMSAs were performed. Twenty µM NTR protein was combined with 10 µM FITC-tagged NLS peptide, in the presence of 7.5% glycerol, and electrophoretically separated on a 1.5% agarose gel in TB buffer (45 mM Tris base, 45 mM boric acid) at 75 V for 1.5-2 hrs. Gels were imaged for FITC peptide detection before staining with Coomassie Blue for protein detection. Protein only and peptide only controls were also run.

### 2.6. Fluorescence polarization (FP) assays

To quantitatively assess the interaction between NLSs and NTRs, FP assays were performed based on previously described methods (39). Twenty µM NTR protein was titrated in a two-fold dilution series across 23 wells of a black Fluotrac microplate (Greiner Bio-One, Kremsmünster, Austria) and combined with 10 nM FITC-tagged NLS peptide. Wells were made up to a total volume of 200 µL with SEC buffer. Fluorescence polarization was measured using a CLARIOstar Plus plate reader (BMG Labtech, Ortenberg, Germany). A peptide only control was included and used for gain adjustment. Assays were performed in triplicate. Data were analyzed in GraphPad Prism (version 10.2.2; GraphPad Software, Boston, MA, USA) using non-linear regression with one site-specific binding to determine the dissociation constant (Kd) and maximum binding (Bmax).

### 2.7. Cell culture

HEK293A cells were maintained in Dulbecco’s modified Eagle’s medium (DMEM) supplemented with 10% (v/v) fetal bovine serum, 50 U/mL penicillin, 50 U/mL streptomycin, and 2 mM L-glutamine (Thermo Fisher Scientific, Monza, Italy), and passaged when confluent (40).

### 2.8. Confocal laser scanning microscopy (CLSM) image analysis

HEK293A cells were seeded onto glass coverslips in a 24-well plate (4 × 10^4^ cells/well) and the next day transfected with appropriate amounts of expression constructs (range 100–250 ng) using Lipofectamine 2000 (Thermo Fisher Scientific, Monza, Italy), as previously described (41). At 48 hrs post-transfection (p.t.), cells were incubated for 30 min with DRAQ5 (1:5000 in DMEM, no phenol red), washed with PHEM 1x (60 mM PIPES, 25 mM HEPES, 10 mM EGTA, 4 mM MgSO_4_), and fixed with 4% (v/v) paraformaldehyde for 10 min. Following three washes with PHEM 1x and one wash with milliQ water, coverslips were mounted on glass slides with Fluoromount G (Southern Biotech, Birmingham, AL, USA). Subcellular localization of fusion proteins was analyzed using a Nikon A1 confocal laser scanning microscope (Nikon, Tokio, Japan) equipped with a 60x oil immersion objective, as described previously (42). DRAQ5 and fibrillarin were used to define nuclear and nucleolar masks, respectively, whereas a small area close to DRAQ5 was used to define a cytosolic mask, as described previously (43). The fluorescence attributed to auto-fluorescence/background (Fb) was subtracted from the measurements to calculate the Fn/c and Fno/Fn ratios according to the formulas Fn/c = (Fn − Fb)/(Fc − Fb) and Fno/Fn = (Fno − Fb)/(Fn − Fb). Cells with oversaturated signals were excluded from analysis. In some cases, to allow easier detection of nucleoli, cells were co-transfected with DsRed-fibrillarin or DsRed-nucleolin expression plasmids and rgb profile plots were calculated with Fiji (44). Statistical analysis was performed using GraphPad Prism (version 9; GraphPad Software, Boston, MA, USA) applying Student’s t test, one-way ANOVA, or two-way ANOVA as appropriate.

### 2.9. Inhibition of Ran-Dependent Nuclear Transport

Ran-dependent nuclear transport was inhibited by depletion of cellular RanGTP resulting from a lack of free GTP (45), by incubating cells for 30 min at 37°C in DMEM containing no glucose, 5% FBS, and supplemented with 10 mmol/L sodium azide, 6 mmol/L 2-deoxy-D-glucose (D8357, Sigma, Merck Millipore, Milan, Italy) and DRAQ5 (#62251; ThermoFisher Scientific, Monza, Italy; 1:5000) as described previously (43, 46), before being stained, fixed and analyzed by CLSM as detailed above.

## 3. Results

### 3.1. TTDoV ORF1 contains putative NLSs located at the N-terminus and the C-terminus

Although TTVs are believed to replicate in the cell nucleus, the ability of TTV encoded proteins to interact with host NTRs has never been investigated. Further, previous studies have reported conflicting results concerning the subcellular localization of ORF1 from different TTV isolates (26-28). To shed some light on this topic, we analyzed the primary amino acid sequence of TTDoV (isolate At-TTV3; Figure 1A) ORF1 with the cNLS Mapper software, searching for putative NLSs. Our analysis revealed that ORF1 possessed several putative NLSs (Figure 1B). In particular, eleven partially overlapping NLSs were predicted in the N-terminal ARM spanning from residues 4 to 75 (the strongest being 27-RRWRRRPRRRRRPYRRRPYRRYGRRRKVRRR-57; NLSn), and one NLS was predicted at the C-terminus (632-LPPPEKRARWGF-643; NLSc). To assess whether the identified NLSs could be accessible to host cell NTRs, AlphaFold3 was used to predict the structure of full length TTDoV ORF1 protein and putative NLSs were mapped onto the predicted model. The model of ORF1 monomers (Figure 1C; Supplementary Table S3) featured largely unstructured N- and C-termini, which is where both putative NLSs were located. Between these regions (residues 71-546) was a single jelly-roll fold comprised of eight β-strands (B-I) forming two anti-parallel β-sheets, the larger BIDG and the smaller CHEF, with a projection domain extending out between β-strands H and I (Supplementary Figure S1), as described for ORF1 proteins from *Betatorquevirus* isolate LY1 and beak and feather disease virus (BFDV), and as predicted for several additional *Anelloviridae* ORF1 proteins (5, 13, 14). Both putative NLSs were modelled in unstructured regions of the protein with low confidence scores (pLDDT < 70); this strongly correlates with regions that are intrinsically disordered (47), thus NLSn and NLSc are proposed to be accessible for binding to NTRs in ORF1 monomers.

**Figure 1.**
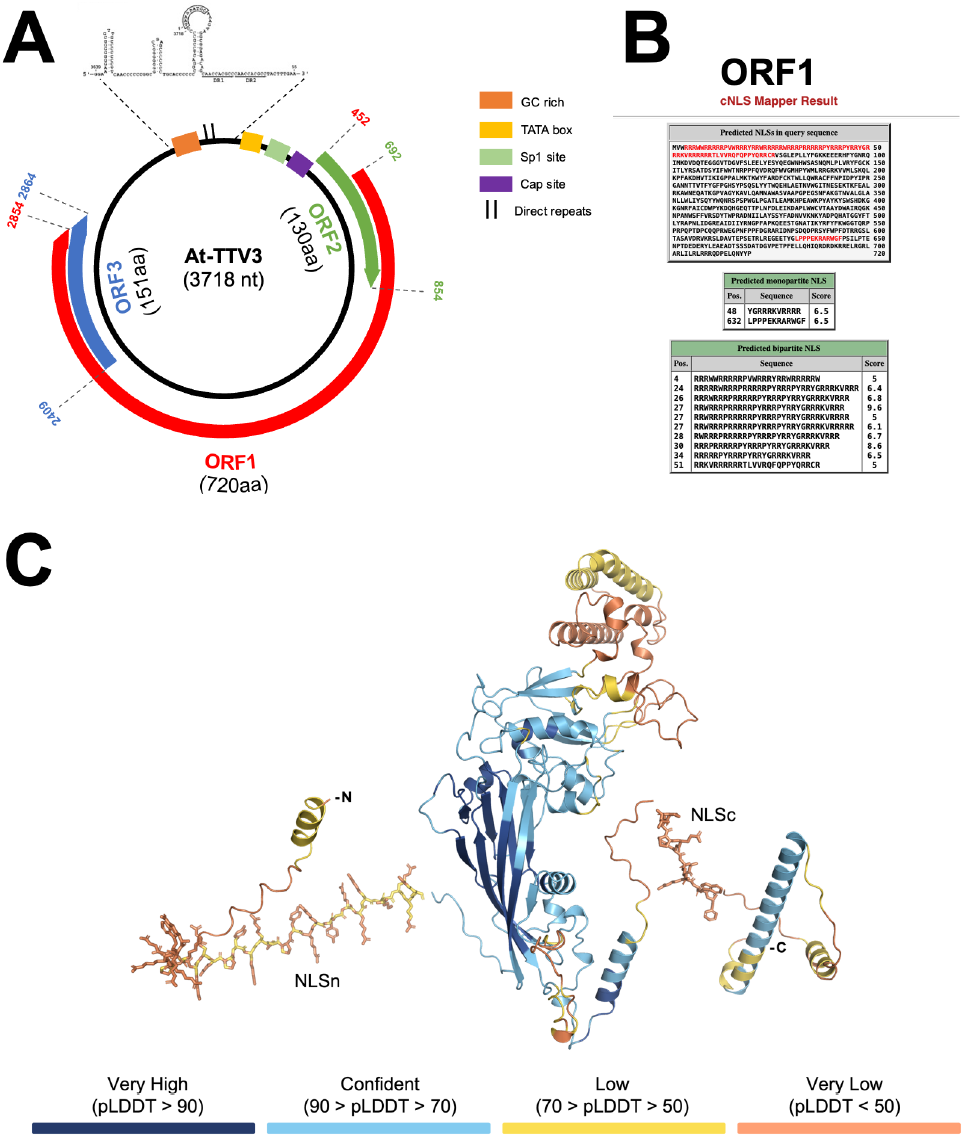
TTDoV ORF1 protein contains putative NLSs that are proposed to be accessible for NTR binding. (A) Schematic representation of TTDoV genome features and ORFs relative to sequence deposited in NCBI. (B) The TTDoV ORF1 amino acid sequence was retrieved from UniProt (UniProt: Q9DUB7) and analyzed with cNLS Mapper for identification of putative NLSs. Top panel: the protein sequence is displayed using the single letter amino acid code, with identified putative NLSs shown in red. Bottom panels: the predicted NLS sequences are shown, along with the position of the first amino acid and the predicted cNLS Mapper score. (C) AlphaFold3 model of TTDoV ORF1 (UniProt: Q9DUB7, residues 1-720). The top ranked prediction is shown; colored by pLDDT score of estimated confidence: very high (pLDDT > 90) in dark blue, confident (90 > pLDDT > 70) in light blue, low (70 > pLDDT > 50) in yellow, and very low (pLDDT < 50) in orange. Model shown in cartoon; putative NLSs shown in stick representation. Both putative NLSs are located in unstructured regions of the protein and are proposed to be accessible for binding to NTRs.

### 3.2. Basic residues within TTDoV ORF1 protein mediate high affinity interactions with several NTRs

We set out to functionally validate the newly identified TTDoV ORF1 putative NLSs by first testing their ability to interact with selected NTRs, including IMPα1/2/3/5/7 and IMPβ1/2/3. Fluorescein isothiocyanate (FITC)-labeled peptides corresponding to the predicted NLS motifs were synthesized and subjected to EMSAs and FP binding assays. While multiple peptides exhibited robust NTR interactions, the peptide corresponding to the NLSn consistently precipitated and failed to migrate during EMSA, suggesting poor solubility under assay conditions. Consequently, we were unable to evaluate binding of the NLSn peptide to any of the tested NTRs using either EMSA or FP, precluding further analysis of this motif (Figure 2A,C-D; Supplementary Figure S2A,C). To investigate further, NLSn was divided into two sections based on cNLS Mapper results, namely the arginine rich region (27-RRWRRRPRRRRRPYRRRPYRR-47; ORF1 NLSn_A) and the predicted monopartite NLS (48-YGRRRKVRRR-57; ORF1 NLSn_B). Both NLSn_A and

**Figure 2.**
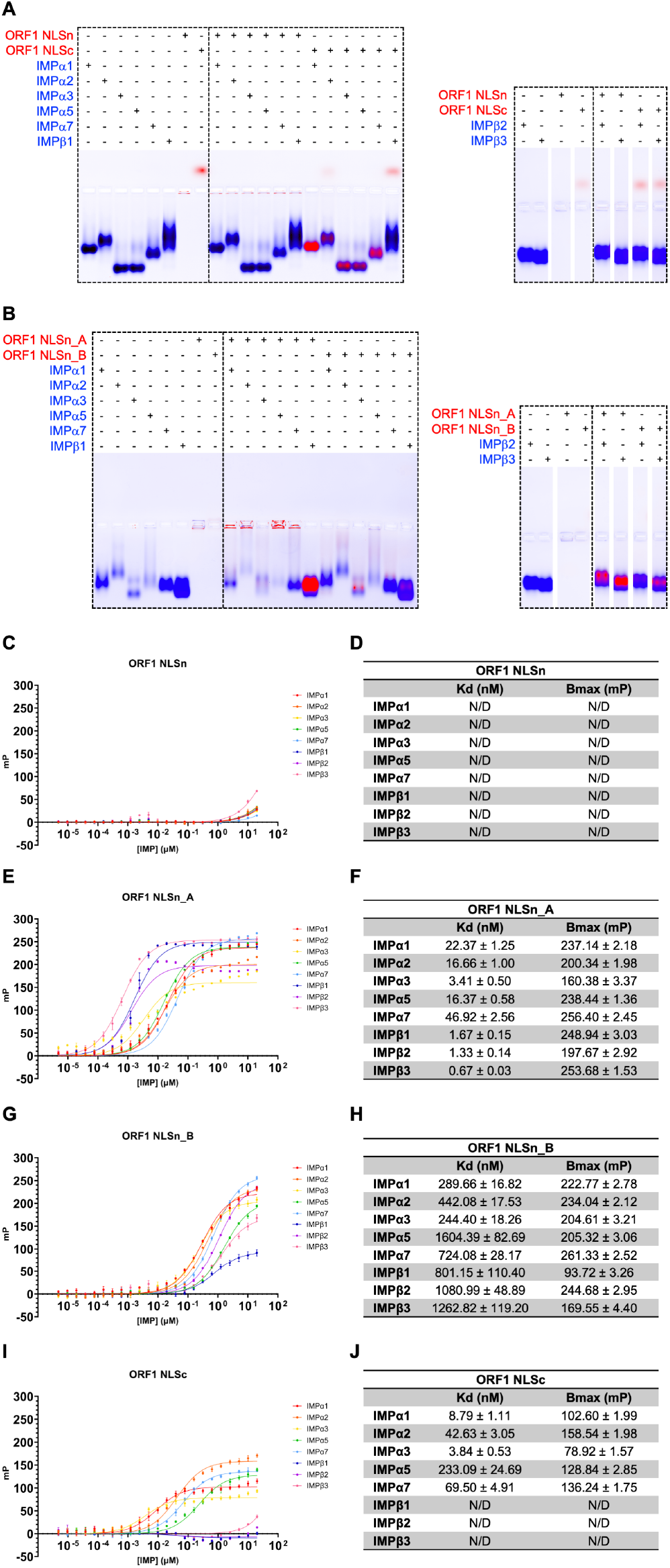
TTDoV ORF1 can bind NTRs through both its N-terminal and C-terminal NLS. (A-B) Electrophoretic mobility shift assays assessing binding of NTRs (20 µM) and FITC-tagged NLS peptides (10 µM). Proteins are shown in blue; peptides are shown in red; co-migration indicates binding; same IMPβ2/3 protein only controls shown in both A and B. (C-J) Fluorescence polarization assays measuring binding affinity between NTRs (20 µM starting concentration) and FITC-tagged NLS peptides (10 nM), including calculated dissociation constant (Kd) and maximum binding (Bmax) values. Data shown as *n* = 3; error bars represent mean ± standard error of the mean; N/D = not determined.

NLSn_B co-migrated with several IMPα and IMPβ NTRs, indicating their potential as NLSs, although some precipitation of the arginine rich NLSn_A peptide was still evident (Figure 2B; Supplementary Figure S2B-C). FP assays confirmed binding of ORF1 NLSn_A and NLSn_B with all tested NTRs. NLSn_A bound all NTRs with high affinity (~1-47 nM), exhibiting a preference for IMPβ’s (all <2 nM) (Figure 2E-F), while ORF1 NLSn_B bound all NTRs with a moderate to low binding affinity (~244-1604 nM), exhibiting a preference for IMPα1 (290 nM) and IMPα3 (244 nM) (Figure 2G-H). ORF1 NLSc bound to all tested IMPα isoforms, with moderate to high binding affinity (~4-233 nM) and a preference for IMPα1 (9 nM) and IMPα3 (4 nM), but not IMPβ’s (Figure 2A,I-J; Supplementary Figure S2A,C). These results suggest that TTDoV ORF1 nuclear import could be mediated by an N-terminal and a C-terminal NLS, capable of binding both IMPα (NLSn and NLSc) and IMPβ (NLSn) NTRs. This raises the possibility that TTDoV ORF1 has evolved to simultaneously exploit multiple nuclear import pathways, as recently described for several cellular (48, 49) and viral (43) proteins.

### 3.2. Basic residues within TTDoV ORF1 protein confer RanGTP- and IMPα/μ1-dependent nuclear targeting properties to heterologous proteins

The observed high affinity of ORF1 NLS peptides for NTRs suggests their involvement in protein nuclear translocation. To investigate this possibility, we measured their ability to confer nuclear targeting properties to GFP. To this end, we quantified the levels of nuclear accumulation of GFP fused to ORF1 NLSs when transiently expressed in mammalian cells, using GFP alone and a GFP-SV40 NLS fusion protein as negative and positive controls, respectively. While GFP evenly distributed throughout the cell, GFP-SV40 NLS strongly accumulated in the cell nucleus (Fn/c of 1.1 and 9.7, respectively; Figure 3). Interestingly, GFP-ORF1 NLSn could be detected both in the nucleus and, to a lesser extent, in the cytoplasm (Fn/c of 3.2; Figure 3), but strongly accumulated in the nucleoli, as evidenced by extensive co-localization with nucleolin (Supplementary Figure S3A) and an Fno/n of 3.0 (Supplementary Figure S3B-C). On the other hand, GFP-ORF1 NLSc was primarily localized in the nucleoplasm, with nucleolar exclusion and an Fn/c of 3.0. Depletion of intracellular RanGTP by incubating cells in media containing sodium azide and 2**-** deoxy-D-glucose (Figure 3B-C, blue circles) or inhibition of the IMPα/β1-dependent nuclear import pathway by overexpression of the competitive IMPα inhibitor Bimax2 (Figure 3B-C, pink circles) resulted in redistribution of all proteins between the nucleus and cytoplasm with a significant loss of nuclear fluorescence, indicating that both NLSs confer active, IMPα/μ1-dependent nuclear transport properties to GFP. However, the nucleolar accumulation of GFP-ORF1 NLSn was unaffected by GTP depletion, thus nucleolar localization mediated by the long stretch of arginine residues is most likely due to interactions with cellular nucleic acids within the nucleoli, rather than an active process.

**Figure 3.**
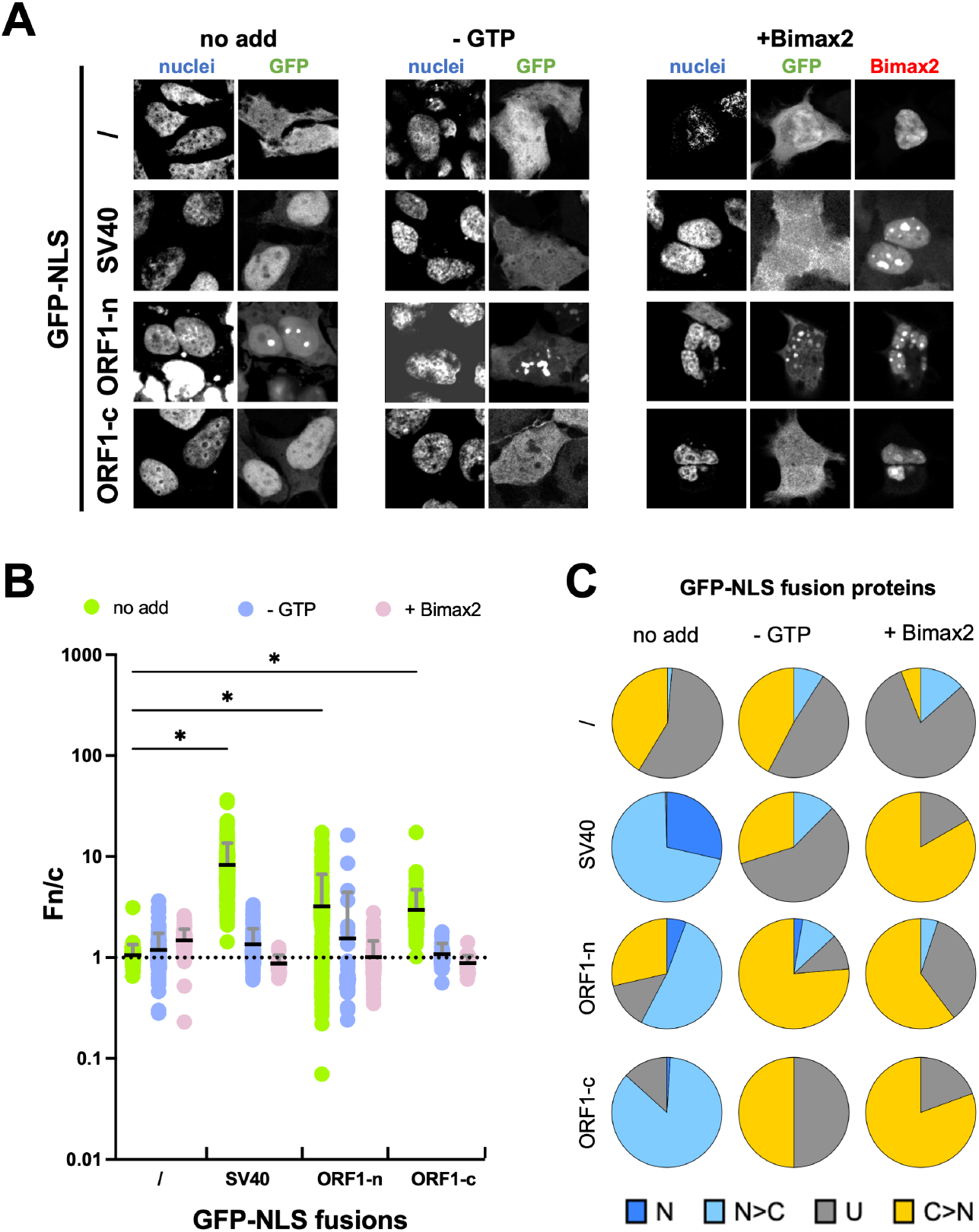
Characterization of nuclear and nucleolar targeting properties of TTDoV ORF1 NLSs in a cellular context. HEK293A cells were transfected to express the indicated GFP fusion proteins, alone or in the presence of mCherry-Bimax2. 24 hrs p.t., cells were either left untreated or incubated for 30 min with an energy depletion media (-GTP) before being stained and fixed for CLSM imaging and analysis. (A) Representative images of the 633 nm (nuclei), 488 nm (GFP), and 561 nm (Bimax2) laser channels are shown, relative to the indicated GFP fusion proteins. (B) Images such as those shown in (A) were analyzed for quantification of the levels of nuclear accumulation (Fn/c) at the single cell level. Data are shown as individual measurements (circles), along with mean (black horizontal bars) and standard deviation of the mean (gray vertical bars), including the results of the Welch and Brown-Forsythe one-way ANOVA for significance between the indicated proteins (*: p ≤ 0.05); pooled data from at least two independent experiments. (C) The percentage of cells relative to each indicated fusion protein displaying the indicated subcellular localization is shown. N: nuclear, Fn/c ≥ 10; N>C: more nuclear than cytosolic, 2 ≤ Fn/c < 10; U: ubiquitous, 1 ≤ Fn/c < 2; C>N: more cytosolic than nuclear, Fn/c < 1.

### 3.4. TTDoV ORF1 nuclear import is primarily dependent on recognition of NLSc by IMPα/β1 while accumulation in the nucleoli relies on the interaction of NLSn with nucleolar components

Our data suggest that ORF1 contains a nucleolar localization signal (NoLS) at residues 27-57 (NLSn) and an NLS at residues 632-643 (NLSc). To more precisely characterize the process of ORF1 intracellular transport and the specific contribution of such sequences to this process, we investigated the subcellular localization of full length ORF1 and derivatives thereof as fused to GFP (Figure 4A). Transient expression of GFP-ORF1 in HEK293A cells resulted in the protein being detected at variable levels in the nucleus and cytosol, with an average Fn/c of 4.0 (Figure 4B-C) and nuclear accumulation in more than 60% of transfected cells (Figure 4D). Further, strong nucleolar accumulation was observed in all cells, highlighted by co-localization with RFP-fibrillarin (Figure 4E) and an average Fno/n of 1.9 (Figure 4F), with nucleolar accumulation in 100% of analyzed cells (Figure 4G). Importantly, nuclear targeting was strongly impaired by overexpression of Bimax2, demonstrating that ORF1 nuclear import is primarily mediated by the IMPα/β1 heterodimer (Figure 4B-D). Deletion of the first 78 amino acids (ORF1 Δ78), containing the ARM and NLSn, resulted in exclusion from the nucleoli (Fno/n of 0.6; Figure 4E-G), but did not reduce nuclear accumulation of ORF1. Rather, the observed redistribution from nucleoli to the nucleoplasm caused a significant increase of the average Fn/c value to 12.4 (Figure 4C). AlphaFold3 models predicted that ORF1 bound IMPα NTRs in the major NLS binding site through NLSc (Figure 4H; Supplementary Table S3), with K637 interacting with key IMPα residues G150, T155, and D192 in the critical P2 binding pocket (Figure 4I). Accordingly, introduction of the K637A substitution resulted in strong impairment of nuclear targeting (average Fn/c of 1.4; Figure 4B-D), but not of nucleolar accumulation (Fno/n of 1.2; Figure 4E-G). Strikingly, introduction of the K637A substitution together with deletion of residues 1-78 resulted in a predominantly cytosolic protein, which failed to accumulate in the nucleus (average Fn/c of 0.8; Figure 4B-D) or the nucleolus (Fno/n of 0.7; Figure 4E-G). Clearly, TTDoV has evolved distinct sequences playing complementary roles in determining ORF1 subcellular localization, with NLSc mediating IMPα/β1-dependent nuclear targeting and NLSn being responsible for nucleolar accumulation, most likely by mediating electrostatic interactions with nucleolar components such as rRNA.

**Figure 4.**
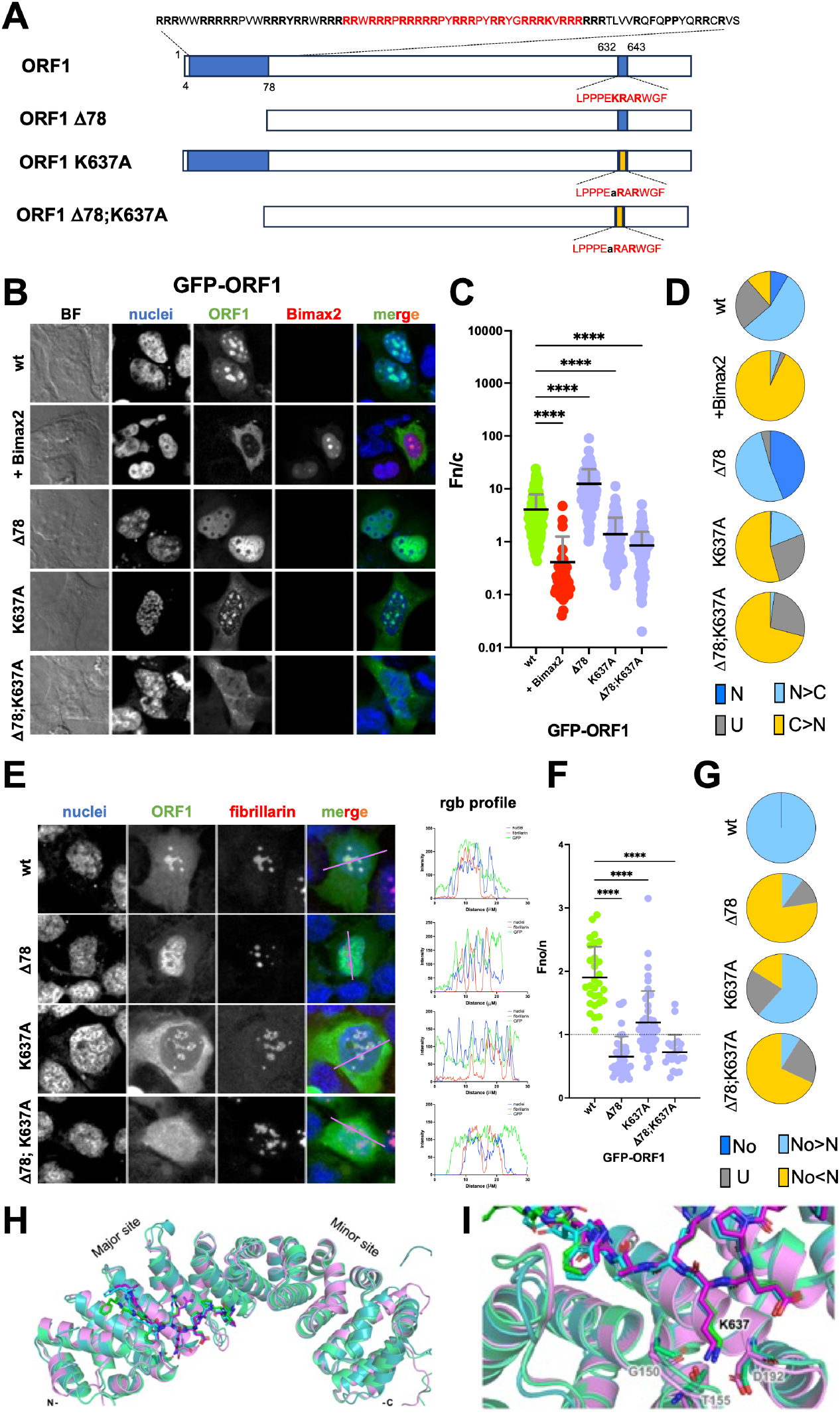
TTDoV ORF1 is translocated into the nucleus by IMPα/β1 which recognizes NLSc, and accumulates in the nucleoli through a NoLS at the N-terminus. (A) Schematic representation of GFP-ORF1 fusion proteins analyzed, along with the respective position and sequence of the targeting signals identified here. ORF1 amino acid sequence is shown as a white box; identified NLS regions are shown as blue boxes; NLS sequences are in red; mutated NLSc is shown as a yellow box. (B) HEK293A cells were transfected with Lipofectamine 2000 to express the indicated GFP-ORF1 fusion proteins in the absence or presence of the IMPα inhibitor Bimax2. Twenty-four hrs p.t., cells were incubated with DRAQ5 to stain cell nuclei, fixed, and processed for CLSM imaging and analysis. Representative images of the bright field (BF), 633 nm (nuclei), 488 nm (GFP), and 561 nm (Bimax2) laser channels are shown, along with a merged image (merge). (C) Images such as those shown in (B) were quantitatively analyzed to calculate the levels of nuclear accumulation (Fn/c) at the single cell level. Data are shown as individual measurements (circles), along with mean (black horizontal bars) and standard deviation of the mean (gray vertical bars), including the results of the Welch and Brown-Forsythe one-way ANOVA for significance between the indicated proteins (****: p ≤ 0.0001); pooled data from at least two independent experiments. (D) The percentage of cells relative to each indicated fusion protein displaying the indicated subcellular localization is shown. N: nuclear, Fn/c ≥ 10; N>C: more nuclear than cytosolic, 2 ≤ Fn/c < 10; U: ubiquitous, 1 ≤ Fn/c < 2; C>N: more cytosolic than nuclear, Fn/c < 1. (E) The indicated GFP fusion proteins were transiently co-expressed in HEK293A cells with DsRed-fibrillarin by means of Lipofectamine 2000 transfection. Twenty-four hrs p.t., cells were incubated with DRAQ5 to stain cell nuclei, fixed, and processed for CLSM imaging and analysis. Representative images of the 633 nm (nuclei), 488 nm (GFP), and 561 nm (fibrillarin) laser channels are shown, along with a merged image (merge) and a rgb profile plot across the indicated area (rgb profile). (F) Images such as those shown in (E) were quantitatively analyzed to calculate the levels of nucleolar accumulation (Fno/n) at the single cell level. Data are shown as individual measurements (circles), along with mean (black horizontal bars) and standard deviation of the mean (gray vertical bars), including the results of the Welch and Brown-Forsythe one-way ANOVA for significance between the indicated proteins (****: p ≤ 0.0001); pooled data from at least two independent experiments. (G) The percentage of cells relative to each indicated fusion protein displaying the indicated subcellular localization is shown. No: nucleolar, Fno/n ≥ 10; No>N: more nucleolar than nuclear, 2 ≤ Fno/n < 10; U: ubiquitous, 1 ≤ Fno/n < 2; No<N: more nuclear than nucleolar, Fno/n < 1. (H) Superposition of AlphaFold3 models of TTDoV ORF1 NLSc bound in the major NLS binding site of IMPα1 (teal), IMPα3 (green), and IMPα7 (purple). (I) K637 is shown to interact with key residues in the P2 binding pocket of IMPα, specifically G150, T155, and D192 (IMPα1 numbering). The top ranked predictions are shown. IMPα shown in cartoon; ORF1 NLSc and IMPα P2 residues shown in stick representation.

### 3.5. Evolution of additional NLSs in capsid proteins from Anelloviridae

The presence of a functional NLS at the C-terminus of TTDoV ORF1, in addition to the one located within the ARM, is noteworthy. In members of the *Circoviridae* family, the capsid protein ARM is known to mediate interactions with host NTRs and is considered the principal determinant of nuclear import (13, 23). Capsid proteins of *Anelloviridae* are thought to have evolved from those of circoviruses through the stepwise acquisition of increasingly complex projection domains within the jelly-roll fold (5, 14). Since TTDoV ORF1 is one of the larget *Anelloviridae* capsid proteins reported so far, and it is predicted to possess a very large projection domain (Supplementary Figure S1) similar to that of alphatorqueviruses (5), we hypothesized that the acquisition of large projection domains could correlate with the presence of additional NLSs downstream of the ARM. To verify this hypothesis, we retrieved the amino acid sequences of capsids from both *Anelloviridae* and *Circoviridae* and scanned them for the presence of putative NLSs (Figure 5, Supplementary Tables S4-5). Intriguingly, NLSs in similar positions were identified in capsids endcoded by several *Anelloviridae* family members, but not in gyroviruses which endcode shorter capsids (Supplementary Figure S4, Supplementary Tables S4-6). Capsids encoded by *Anelloviridae* are significantly longer and more variable in length than those encoded by *Circoviridae* (592±108 versus 295±27 amino acids, respectively; Figure 5A), consistent with the acquisition of projection domains of increasing size (5). Furthermore, the number (Figure 5B) and predicted activity (Figure 5C) of NLSs is higher in anelloviruses than in circoviruses. Moreover, in circoviruses the NLS is most frequently located within the ARM, while more than 70% of anelloviruses possess an additional NLS downstream (Figure 5D). Linear regression showed significant positive correlation between capsid length and NLS activity among anelloviruses (Figure 5E) but not circoviruses (Figure 5F). Similar results were obtained by analyzing the correlation between capsid length and cNLS number (Figure 5G) and cNLS score (Figure 5H) among the most abundant *Anelloviridae* genera. In addition, the percentage of capsids bearing an additional NLS increased progressively with capsid size (Figure 5I). Such NLSs were never identified in gyrovirus capsids, which are significantly shorter than those from other genera (432.9±46.7 amino acids), while they were systematically found in all alphatorqueviruses tested, which encode for the largest capsids (732.4±41.0 amino acids). Therefore, the acquisition of an additional cNLS downstream of the ARM represents a hallmark of *Anelloviridae* evolution and correlates with the presence of a large projection domain.

**Figure 5.**
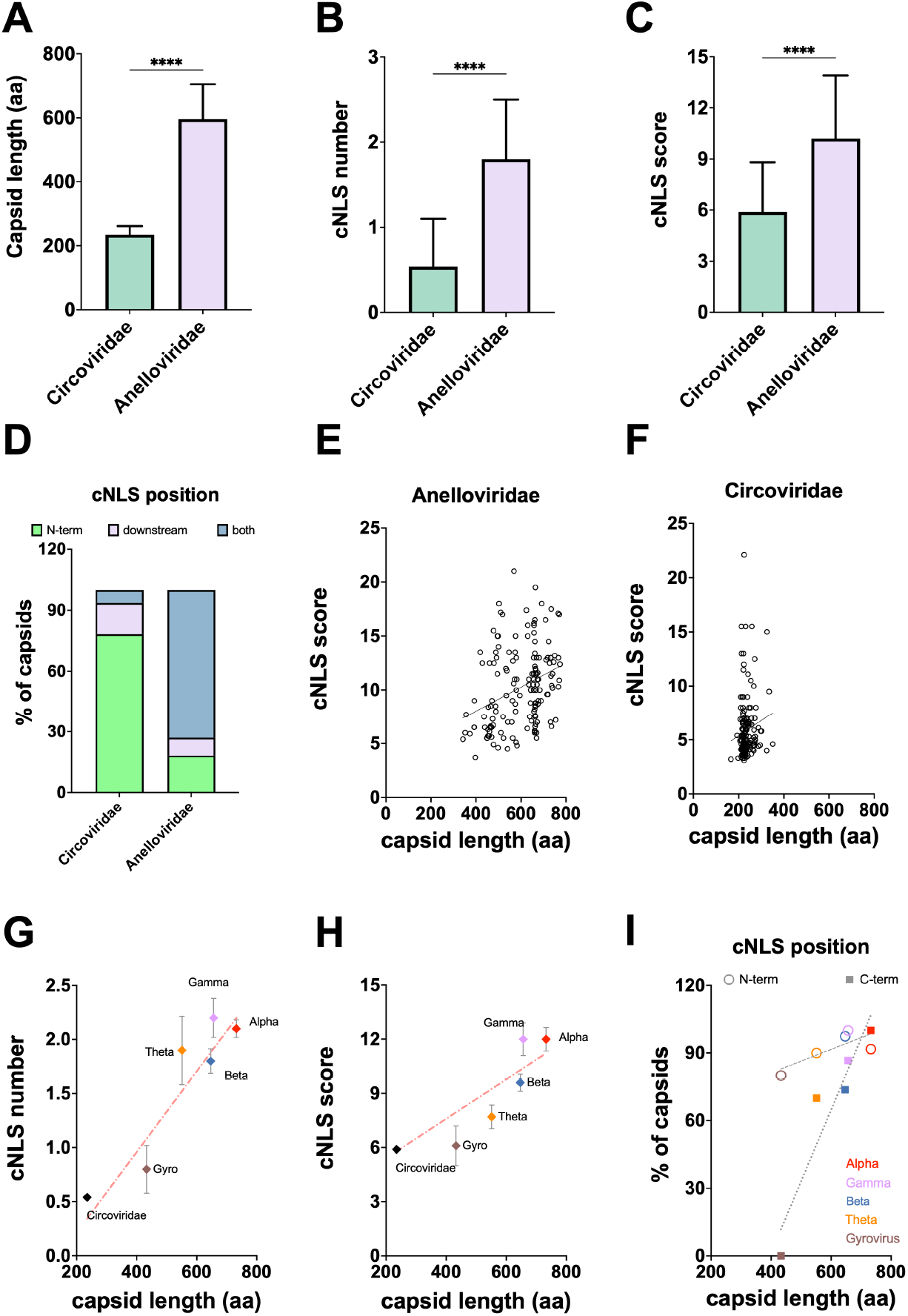
Evolution of C-terminal cNLSs downstream of the ARM correlates with capsid size and represents a potential evolutionary hallmark of *Anelloviridae*. Capsid protein sequences from members of the *Circoviridae* and *Anelloviridae* families were retrieved from UniProt and analyzed using cNLS Mapper to identify putative classical nuclear localization signals (cNLSs) with a predicted score ≥ 5. (A-C) Average capsid protein length (A), number of predicted cNLSs (B), and predicted cNLS activity score (C) for *Circoviridae* and *Anelloviridae*. Data are presented as mean ± standard deviation. Statistical significance was assessed using two-way ANOVA (****: p ≤ 0.0001). (D) Proportion of capsid proteins bearing a predicted cNLS exclusively within the ARM (N-term), exclusively downstream of the ARM (downstream), or in both positions (both). (E-F) Linear regression between capsid length and cNLS activity score in *Anelloviridae* (E) and *Circoviridae* (F). (G-H) Linear regression between capsid length and cNLS number (G) and cNLS activity score (H) across *Circoviridae* and the indicated anellovirus genera. (I) Linear regression between capsid length and the percentage of capsid proteins containing a cNLS within the ARM (N-term) or downstream (C-term) across *Circoviridae* and the indicated anellovirus genera.

## 4. Discussion

The landscape of *Anelloviridae*-host interaction is largely uncharacterized, mainly due to the lack of suitable cellular systems to study the virus life cycle. In this context, little is known regarding the functional interaction between host cell NTRs and capsid proteins. Molecular studies of the virus-host interface may shed light on their potential pathogenicity. Keeping this in mind, we used TTDoV as a model to study the interaction between TTVs and the host cell nuclear transport machinery. Our study identified distinct signals responsible for ORF1 nuclear import and nucleolar targeting, as well as the NTRs responsible for this process. This is the first study to investigate the physical and functional consequences of the interaction between TTV proteins and the host cell nuclear transport machinery, paving the way to a better understanding of TTV-host cell interaction, pathogenicity, and evolution.

### Role of TTDoV ORF1 ARM in nucleolar accumulation

The capsid proteins from several DNA and RNA viruses, including TTV1 P/1C1, are known to localize in the nucleolus to modulate host transcription, process RNA, and promote viral replication (27, 50-56). When expressed in the absence of other viral proteins, TTDoV ORF1 strongly accumulated in the nucleolus (Figure 4B, wt). Such nucleolar accumulation depends on an N-terminal NLSn located within the ARM. Although the ARM has been implicated in genome packaging and nuclear targeting for closely related *Circoviridae* capsids (13, 23), our data indicate that in the context of TTDoV, the ARM primarily supports genome packaging and nucleic acid binding rather than nuclear import. Firstly, TTDoV ORF1 NLSn is sufficient to confer strong nucleolar localization to GFP (Figure 3, ORF1-n; Supplementary Figure S3) and is absolutely required for nucleolar—but not nuclear—targeting of the full-length protein, as evidenced by the strong nucleoplasmic localization observed upon deletion of the first 78 amino acids (Figure 4B-D, Δ78). Secondly, while TTDoV ORF1 nuclear import strongly relies on IMPα/β1, nucleolar accumulation is independent of this pathway (Figure 4). Taken together, these results suggest that nucleolar localization is dependent upon the ability of the arginine-rich NLSn to interact with nucleic acids, which are highly abundant in the nucleolus, or other nucleolar components, as demonstrated for capsid proteins from other DNA and RNA viruses (50-52).

### Role of TTDoV ORF1 NLSc in IMPα/β1-dependent nuclear import

Although TTDoV ORF1 NLSn can bind multiple IMPα and IMPβ NTRs with high affinity (Figure 2B,E-H), ORF1 nuclear import primarily relies on the downstream NLSc, which preferentially interacts with IMPα’s (Figure 2A,I-J) and is predicted to bind the major site of IMPα (Figure 4H). The K637A substitution, targeting a predicted key binding determinant within NLSc (Figure 4I), significantly decreased nuclear levels of ORF1 (Figure 4B-D, K637A). Simultaneous deletion of NLSn further reduced nuclear targeting (Figure 4B-D, Δ78;K637A), indicating that while NLSc is the main driver of ORF1 nuclear import, NLSn also contributes to the process. Interestingly, although Bimax2-mediated inhibition of the IMPα/β1-dependent nuclear import pathway almost completely blocked nuclear localization of ORF1, some protein still accumulated in the nucleolus (Figure 4B-D, +Bimax2). This observation is consistent with the ability of ORF1 residues 27-RRWRRRPRRRRRPYRRRPYRR-47 to mediate high-affinity interactions with IMPβ1, IMPβ2, and IMPβ3 (Figure 2E-F).

### Acquisition of additional NLSs as a hallmark of Anelloviridae capsid evolution

This redundancy of targeting signals and NTR specificity is not unusual and has previously been reported for other viral proteins, including pVII from both human and animal adenoviruses (43), however it does appear to distinguish *Anelloviridae* from *Circoviridae* capsid proteins. Indeed, *Anelloviridae* capsids are thought to have evolved from that of *Circoviridae* through the progressive acquisition of increasingly large projection domains between β-strands H-I of the jelly-roll fold (5, 13, 14). Like alphatorqueviruses (5), TTDoV ORF1 possesses a remarkably large projection domain (Supplementary Figure S1). Our analyses showed that longer projection domains correlate with the acquisition of additional NLSs located downstream of the ARM (Figure 5I). In this respect, gyroviruses appear very similar to *Circoviridae* in that they lack projection domains and contain only an N-terminal NLS within the ARM. In contrast, all alphatorqueviruses possess large projection domains and encode an additional downstream NLS (Figure 5I; Supplementary Figure S4). Intriguingly, the capsid of the *Circoviridae* BFDV lacks a C-terminal NLS. This protein can form distinct macromolecular assemblies depending on the presence of ssDNA, forming large 60-mer VLPs in the presence of DNA and smaller 10-mer VLPs in its absence. The ARM is positioned on the interior of the capsid interacting with DNA in the large VLPs, but remains exposed and accessible for NTR binding in the smaller ones (13, 23, 57). It is therefore plausible that the acquisition of larger projection domains limits the formation of these smaller capsid assemblies where the ARM is exposed, necessitating the evolution of an additional NLS downstream. This hypothesis is supported by the cryo-EM structure of the LY1 torque teno mini virus capsid protein that forms only 60-mer icosahedral capsids, with the ARM buried within the interior (14). Superposition of the predicted TTDoV ORF1 model onto this structure revealed that ORF1 NLSn and NLSc are likely located within the interior and exterior of the viral capsid, respectively (Supplementary Figure S5).

### A model for TTDoV capsid nuclear transport

Our data suggest that the two NLSs play different roles during viral entry and assembly. During viral replication, ORF1 synthesized in the cytoplasm could be translocated into the nucleus by the concerted action of NLSn and NLSc, predominantly through IMPα/β1 and supported by IMPβ1, IMPβ2, and IMPβ3. Subsequently, NLSn located within the ARM could bind viral DNA during packaging and assembly. As a consequence, NLSn would not be available for interaction with NTRs during the first steps of new rounds of viral infection, thus nuclear import of the viral genome is likely mediated by IMPα/β1 through recognition of NLSc. In this context, the acquisition of an additional downstream NLS might represent a particular evolutionary path pursued by *Anelloviridae* members to maximize nuclear import of viral capsid proteins.

Overall, by elucidating the functional interplay between TTDoV ORF1 and host NTRs, our study provides critical mechanistic insight into TTV biology and a conceptual framework for understanding the evolution of nuclear trafficking strategies in small DNA viruses.

## Supplementary Materials

The following supporting information can be found online: Supplementary Figure S1: TTDoV ORF1 contains a jelly-roll fold shared with other capsid proteins; Supplementary Figure S2: TTDoV ORF1 putative NLSs interact with multiple NTRs; Supplementary Figure S3: ORF1 NLSn confers nucleolar targeting properties to GFP; Supplementary Figure S4: Evolutionary relationship between *Anelloviridae* capsids and acquisition of additional NLSs; Supplementary Figure S5: Internalization of the TTDoV ORF1 N-terminus likely renders NLSn inaccessible for binding; Supplementary Table S1: List of plasmids used in this study; Supplementary Table S2: List of peptides used in this study; Supplementary Table S3: Input parameters and confidence metrics of the top five AlphaFold3 predictions; Supplementary Table S4: Analysis of cNLSs within capsid proteins from *Anelloviridae* members; Supplementary Table S5: Analysis of cNLSs within capsid proteins from *Circoviridae* members; Supplementary Table S6: NLSc is highly conserved in *Anelloviridae* capsid proteins, depending on protein length.

## Author Contributions

Conceptualization, G.A.; methodology, D.A. and O.T.; validation, G.F.P. and G.A.; formal analysis, G.F.P., S.P., S.N. and G.A.; investigation, G.F.P., S.P. and S.N.; resources, D.A. and O.T.; data curation, G.F.P. and G.A.; writing—original draft preparation, G.F.P. and G.A.; writing—review and editing, G.F.P., S.P., S.N., S.S. and G.A.; visualization, G.F.P., S.P. and G.A.; supervision, J.K.F. and G.A.; project administration, G.A.; funding acquisition, J.K.F. and G.A. All authors have read and agreed to the published version of the manuscript.

## Institutional Review Board Statement

Not applicable.

## Informed Consent Statement

Not applicable.

## Data Availability Statement

The original data presented in the study are openly available in Research data UNIPD at https://researchdata.cab.unipd.it/ (to be filled in during proofs).

## Acknowledgments

We thank Yoshihiro Yoneda, Masahiro Oka, and Denis Archambault for sharing expression plasmids. We thank Jeff Nanson for advice with structural modeling and constructive criticism of the manuscript.

