## Supplementary Figures S1-S5; Supplementary Tables S1, S2, S3, S6 for "Nuclear trafficking of *Anelloviridae* capsid protein ORF1 reflects modular evolution of subcellular targeting signals"

### Supplementary Figure S1

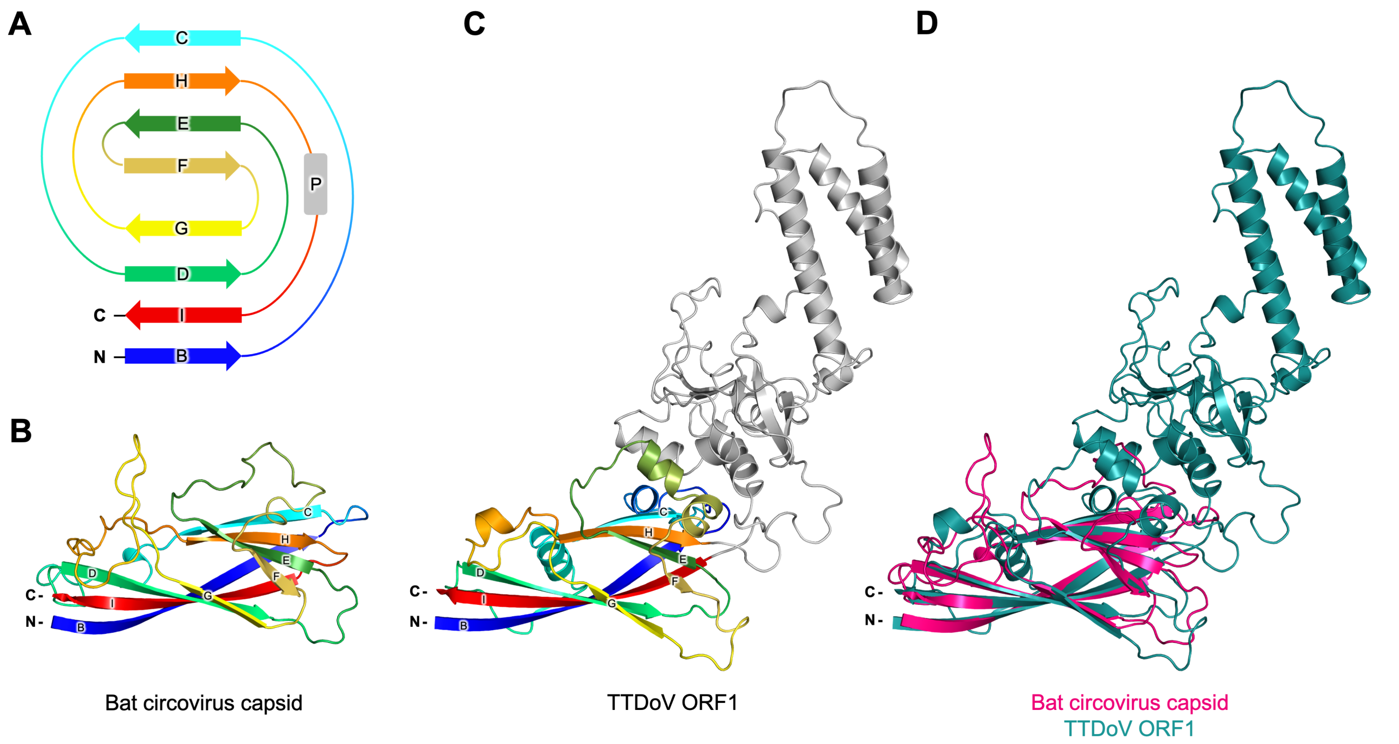


**<u>Supplementary Figure S1.</u> TTDoV ORF1 contains a jelly-roll fold shared with other capsid proteins.** (A) Schematic of jelly-roll fold topology. Arrows represent β-strands labelled B-I, that form two anti-parallel β-sheets (BIDG and CHEF); colored in rainbow from N-terminus (blue) to C-terminus (red). Gray rectangle represents the location of the projection (P) domain found in viral capsid proteins. (B) The jelly-roll fold of the bat *Circovirus* capsid protein (PDB ID: 6RPO) (residues 43-223). Colored as in A. (C) The predicted jelly-roll fold of TTDoV ORF1 (residues 71-546; largely unstructured N- and C-termini containing NLSn and NLSc not shown). Colored as in A. (D) Superposition of the jelly-roll folds of bat *Circovirus* capsid protein (pink) and TTDoV ORF1 (teal), with an RMSD of 3.48 Å over 71 residues (Cα atoms).

### Supplementary Figure S2

A

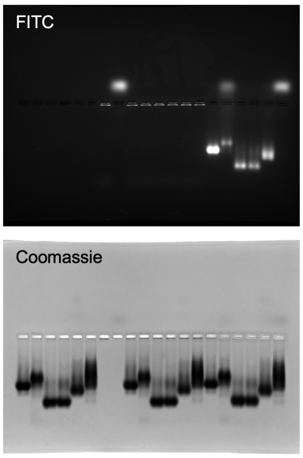


| Lane | Sample |
|---|---|
| 1 | IMPα1 |
| 2 | IMPα2 |
| 3 | IMPα3 |
| 4 | IMPα5 |
| 5 | IMPα7 |
| 6 | IMPβ1 |
| 7 | ORF1 NLSn |
| 8 | ORF1 NLSc |
| 9 | IMPα1 + ORF1 NLSn |
| 10 | IMPα2 + ORF1 NLSn |
| 11 | IMPα3 + ORF1 NLSn |
| 12 | IMPα5 + ORF1 NLSn |
| 13 | IMPα7 + ORF1 NLSn |
| 14 | IMPβ1 + ORF1 NLSn |
| 15 | IMPα1 + ORF1 NLSc |
| 16 | IMPα2 + ORF1 NLSc |
| 17 | IMPα3 + ORF1 NLSc |
| 18 | IMPα5 + ORF1 NLSc |
| 19 | IMPα7 + ORF1 NLSc |
| 20 | IMPβ1 + ORF1 NLSc |

B

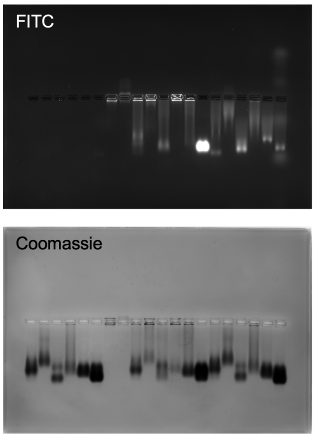


| Lane | Sample |
|---|---|
| 1 | IMPα1 |
| 2 | IMPα2 |
| 3 | IMPα3 |
| 4 | IMPα5 |
| 5 | IMPα7 |
| 6 | IMPβ1 |
| 7 | ORF1 NLSn_A |
| 8 | ORF1 NLSn_B |
| 9 | IMPα1 + ORF1 NLSn_A |
| 10 | IMPα2 + ORF1 NLSn_A |
| 11 | IMPα3 + ORF1 NLSn_A |
| 12 | IMPα5 + ORF1 NLSn_A |
| 13 | IMPα7 + ORF1 NLSn_A |
| 14 | IMPβ1 + ORF1 NLSn_A |
| 15 | IMPα1 + ORF1 NLSn_B |
| 16 | IMPα2 + ORF1 NLSn_B |
| 17 | IMPα3 + ORF1 NLSn_B |
| 18 | IMPα5 + ORF1 NLSn_B |
| 19 | IMPα7 + ORF1 NLSn_B |
| 20 | IMPβ1 + ORF1 NLSn_B |

C

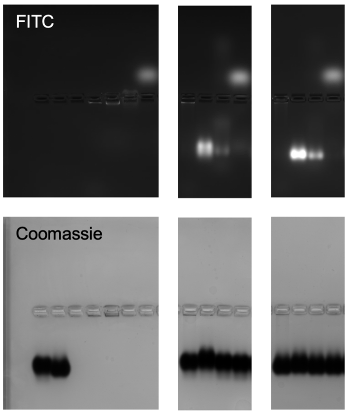


| Lane | Sample |
|---|---|
| 1 | IMPβ2 |
| 2 | IMPβ3 |
| 3 | - |
| 4 | ORF1 NLSn |
| 5 | ORF1 NLSn_A |
| 6 | ORF1 NLSn_B |
| 7 | ORF1 NLSc |
| 8 | IMPβ2 + ORF1 NLSn |
| 9 | IMPβ2 + ORF1 NLSn_A |
| 10 | IMPβ2 + ORF1 NLSn_B |
| 11 | IMPβ2 + ORF1 NLSc |
| 12 | IMPβ3 + ORF1 NLSn |
| 13 | IMPβ3 + ORF1 NLSn_A |
| 14 | IMPβ3 + ORF1 NLSn_B |
| 15 | IMPβ3 + ORF1 NLSc |

<u>Supplementary Figure S2.</u> **TTDoV ORF1 putative NLSs interact with multiple NTRs.** Electrophoretic mobility shift assays assessing binding of NTRs (20 µM) and FITC-tagged NLS peptides (10 µM). Left: Top panel shows FITC expression of peptides; bottom panel shows Coomassie Blue staining of proteins. Right: Table shows samples run in each lane. (A) ORF1 NLSn and ORF1 NLSc peptides against IMPα isoforms and IMPβ1. (B) ORF1 NLSn_A and ORF1 NLSn_B peptides against IMPα isoforms and IMPβ1. (C) IMPβ2 and IMPβ3 against ORF1 NLSn, ORF1 NLSn_A, ORF1 NLSn_B, and ORF1 NLSc peptides.

### Supplementary Figure S3

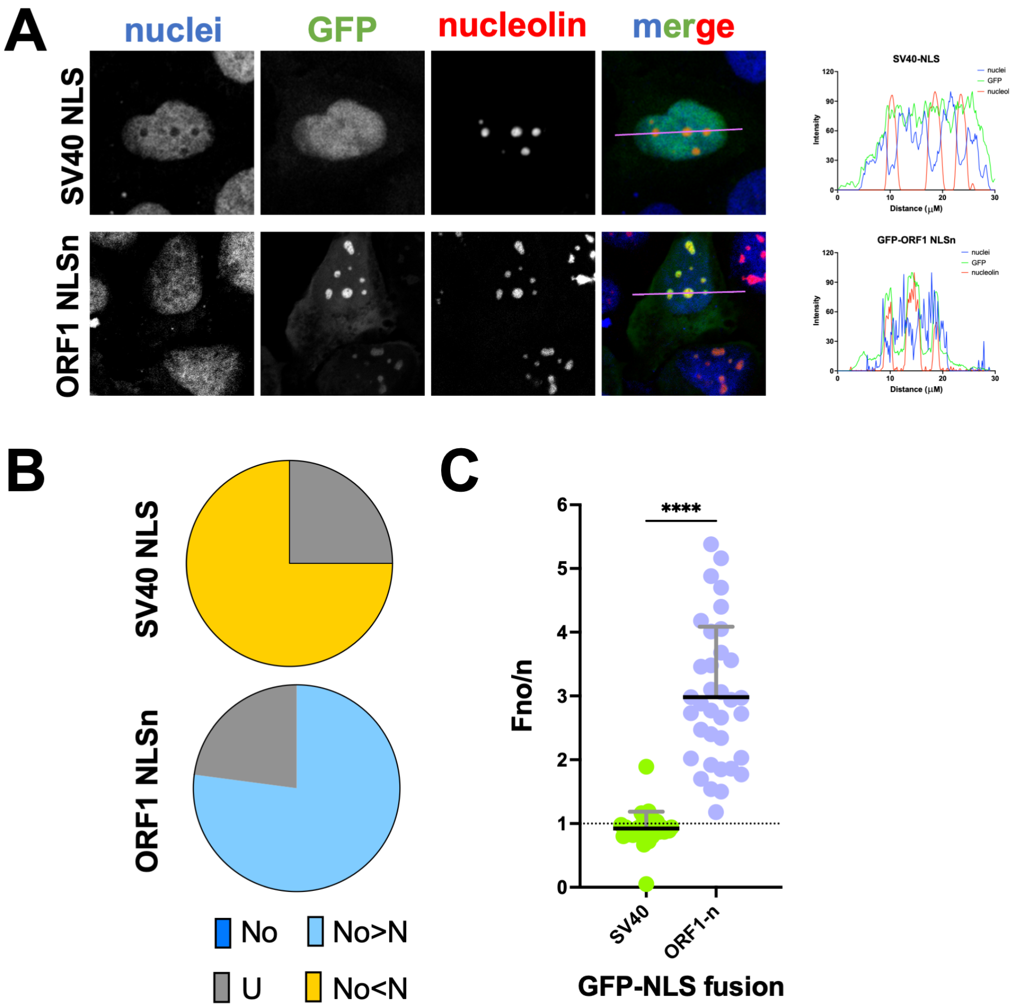


__Supplementary Figure S3.__ **ORF1 NLSn confers nucleolar targeting properties to GFP.** (A) The indicated GFP fusion proteins were transiently co-expressed in HEK293A cells with DsRed-nucleolin by means of Lipofectamine 2000 transfection. Twenty-four hrs p.t., cells were incubated with DRAQ5 to stain cell nuclei, fixed, and processed for CLSM imaging and analysis. Representative images of the 633 nm (nuclei), 488 nm (GFP), and 561 nm (nucleolin) laser channels are shown, along with a merged image (merge) and a rgb profile plot across the indicated area. (B) Images such as those shown in (A) were quantitatively analyzed to calculate the levels of nucleolar accumulation (Fno/n) at the single cell level. The percentage of cells relative to each indicated fusion protein displaying the indicated subcellular localization is shown. No: nucleolar, Fno/n $\geq$ 10; No>N: more nucleolar than nuclear, 2 $\leq$ Fno/n < 10; U: ubiquitous, 1 $\leq$ Fno/n < 2; No<N: more nuclear than nucleolar, Fno/n < 1. (C) Data are shown as individual measurements (circles), along with mean (black horizontal bars) and standard deviation of the mean (gray vertical bars), including the results of the student's t test (****: p ≤ 0.0001); pooled data from two independent experiments.

### Supplementary Figure S4

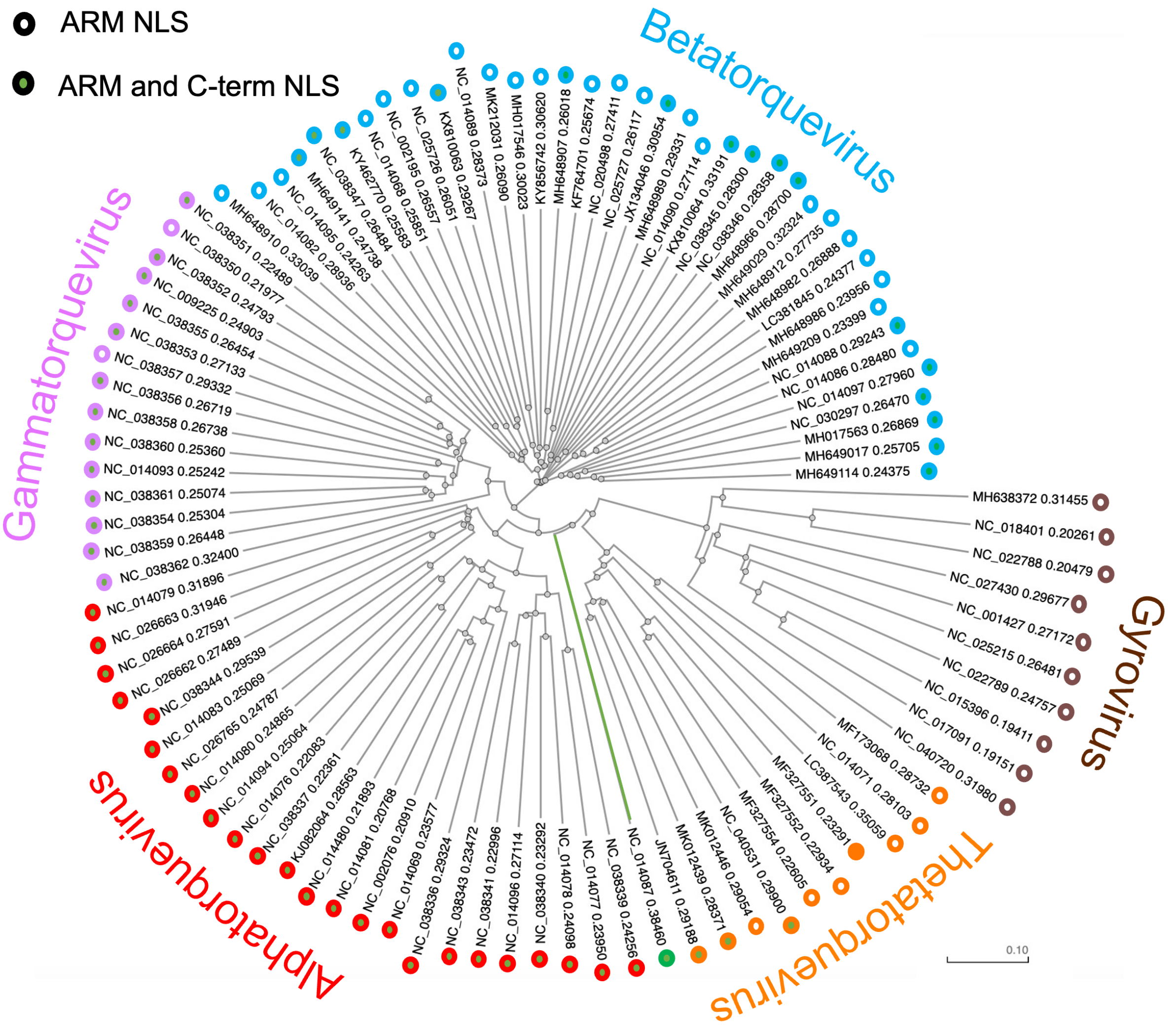


**<u>Supplementary Figure S4.</u> Evolutionary relationship between *Anelloviridae* capsids and acquisition of additional NLSs.** The sequence of ORF1 proteins from members of the indicated *Anelloviridae* genre were phylogenetically analyzed using Clustal Omega. Each protein is indicated with either an empty circle or a circle filled in green, depending on the absence or presence of an NLS downstream of the ARM, respectively. Circle borders are colored according to the viral genera as indicated, while TTDoV ORF1 is colored green.

### Supplementary Figure S5

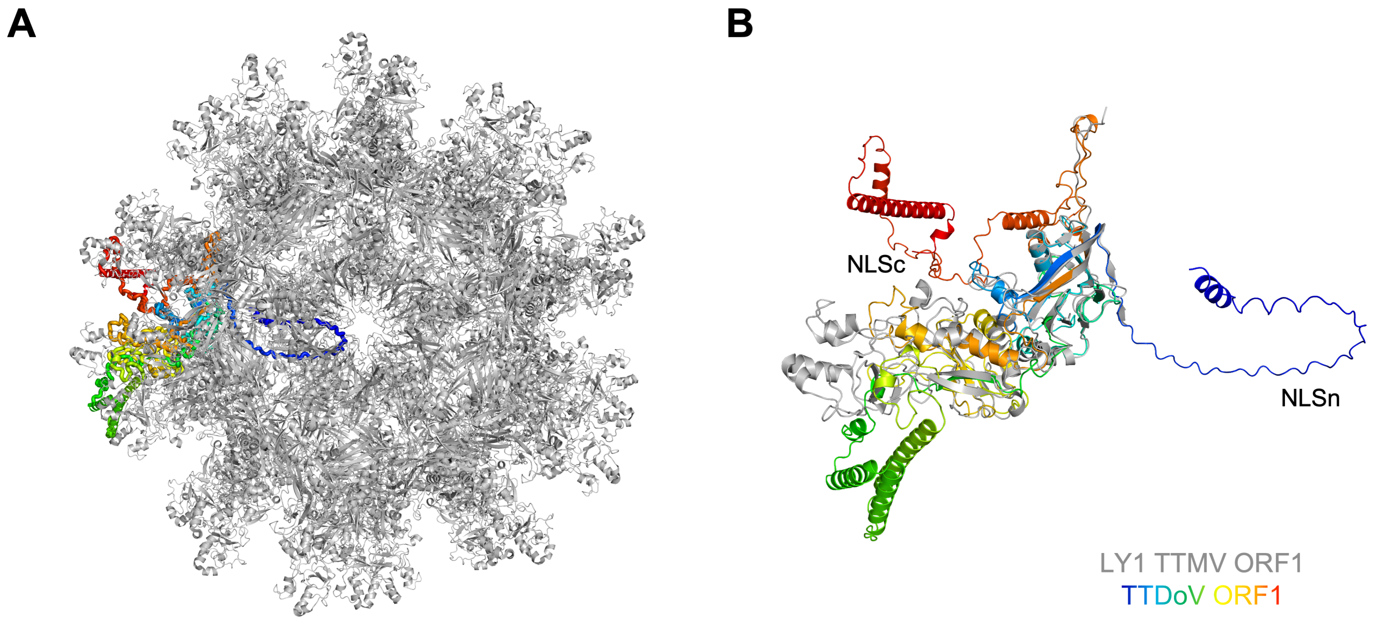


**<u>Supplementary Figure S5.</u> Internalization of the TTDoV ORF1 N-terminus likely renders NLSn inaccessible for binding.** Superposition of the predicted TTDoV ORF1 model onto the cryo-EM structure of LY1 torque teno mini virus (TTMV) ORF1 (PDB ID: 8V7X), with an RMSD of 1.33 Å over 273 residues (Cα atoms). LY1 TTMV ORF1 forms a 60-mer assembly arranged with icosahedral *T*=1 symmetry. If TTDoV ORF1 assembles in a similar manner, the N-terminus (containing NLSn) would be buried within the interior of the virus, and the C-terminus (containing NLSc) would be accessible on the exterior of the virus. LY1 TTMV ORF1 colored in gray; TTDoV ORF1 colored in rainbow from N-terminus (blue) to C-terminus (red). (A) 60-mer. (B) Monomer.

**Supplementary Table S1. List of plasmids used in this study.**

| Plasmid name | Expression product | Source |
|---|---|---|
| hIMPα1ΔIBB_pET-30a(+) | Human importin alpha 1 (*KPNA2*); UniProt: P52292; residues 71-529 | GenScript |
| mIMPα2ΔIBB_pET-30a(+) | Mouse importin alpha 1 (*Kpna2*); UniProt: P52293; residues 70-529 | (58) |
| hIMPα3ΔIBB_pET-30a(+) | Human importin alpha 3 (*KPNA4*); UniProt: O00629; residues 64-521 | GenScript |
| hIMPα5ΔIBB_pET-30a(+) | Human importin alpha 5 (*KPNA1*); UniProt: P52294; residues 74-538 | GenScript |
| hIMPα7ΔIBB_pET-30a(+) | Human importin alpha 7 (*KPNA6*); UniProt: O60684; residues 74-536 | GenScript |
| hIMPβ1_pMCSG21 | Human importin beta 1 (*KPNB1*); UniProt: Q14974; residues 1-876 | This study |
| hIMPβ2_pET-30a(+) | Human importin beta 2 (*TNPO1*); UniProt: Q92973; residues 1-890 | GenScript |
| hIMPβ3_pET-30a(+) | Human importin beta 3 (*IPO5*); UniProt: O00410; residues 1-1097 | GenScript |
| pCDNA3-mCherry-Bimax2 | mCherry-Bimax2 | (59) |
| pNT-TOPO-GFP | GFP | (33) |
| pNT-TOPO-GFP-SV40-LTA-NLS | GFP-SV40 LTA (126-PKKKRKV-132) | (33) |
| pDsRed-C1-fibrillarin | DsRed-fibrillarin | (35) |
| pDsRed-C1-nucleolin | DsRed-nucleolin | (35) |
| pNT-TOPO-GFP-ORF1-NLSn | GFP-ORF1-NLSn (27-RRWRRRPRRRRRPYRRRPYRRYGRRKVRRR-57) | This study |
| pNT-TOPO-GFP-ORF1-NLSc | GFP-ORF1-NLSc (632-LPPPEKRARWGF-643) | This study |
| pEGFP-C1-ORF1 | GFP-ORF1 | This study |
| pEGFP-C1-ORF1Δ78 | GFP-ORF1 lacking residues 1-78 | This study |
| pEGFP-C1-ORF1;K637A | GFP-ORF1 bearing the K637A substitution | This study |
| pEGFP-C1-ORF1Δ78;K637A | GFP-ORF1 lacking residues 1-78 and bearing the K637A substitution | This study |

**Supplementary Table S2. List of peptides used in this study.**

| Peptide name | Peptide sequence | Modification | Source |
|---|---|---|---|
| ORF1 NLSn | 27-RRWRRRPRRRRRPYRRRPYRRYGRRKVRRR-57 | N-terminal FITC-AhX | This study |
| ORF1 NLSn_A | 27-RRWRRRPRRRRRPYRRRPYRR-47 | N-terminal FITC-AhX | GenScript |
| ORF1 NLSn_B | 48-YGRRRKVRRR-57 | N-terminal FITC-AhX | GenScript |
| ORF1 NLSc | 632-LPPPEKRARWGF-643 | N-terminal FITC-AhX | This study |

**Supplementary Table S3. Input parameters and confidence metrics of the top five AlphaFold3 predictions.**

| | pTM* | ipTM^ |
|---|---|---|
| **TTDoV ORF1** | | |
| **Input:** UniProt: Q9DUB7; residues 1-720 (1 copy) | | |
| AlphaFold3_Model_TTDoV_ORF1_0 | 0.55 | N/A |
| AlphaFold3_Model_TTDoV_ORF1_1 | 0.55 | N/A |
| AlphaFold3_Model_TTDoV_ORF1_2 | 0.55 | N/A |
| AlphaFold3_Model_TTDoV_ORF1_3 | 0.55 | N/A |
| AlphaFold3_Model_TTDoV_ORF1_4 | 0.55 | N/A |
| **TTDoV ORF1: IMPα1** | | |
| **Input:** UniProt: Q9DUB7; residues 1-720 (1 copy) + UniProt: P52292; residues 71-529 (1 copy) | | |
| AlphaFold3_Model_TTDoV_ORF1-IMPα1_0 | 0.44 | 0.60 |
| AlphaFold3_Model_TTDoV_ORF1-IMPα1_1 | 0.44 | 0.62 |
| AlphaFold3_Model_TTDoV_ORF1-IMPα1_2 | 0.45 | 0.61 |
| AlphaFold3_Model_TTDoV_ORF1-IMPα1_3 | 0.44 | 0.62 |
| AlphaFold3_Model_TTDoV_ORF1-IMPα1_4 | 0.44 | 0.61 |
| **TTDoV ORF1: IMPα3** | | |
| **Input:** UniProt: Q9DUB7; residues 1-720 (1 copy) + UniProt: O00629; residues 64-521 (1 copy) | | |
| AlphaFold3_Model_TTDoV_ORF1-IMPα3_0 | 0.44 | 0.61 |
| AlphaFold3_Model_TTDoV_ORF1-IMPα3_1 | 0.44 | 0.61 |
| AlphaFold3_Model_TTDoV_ORF1-IMPα3_2 | 0.44 | 0.59 |
| AlphaFold3_Model_TTDoV_ORF1-IMPα3_3 | 0.44 | 0.61 |
| AlphaFold3_Model_TTDoV_ORF1-IMPα3_4 | 0.43 | 0.17 |

| | | |
|---|---|---|
| **TTDoV ORF1: IMPα7** | | |
| **Input:** UniProt: Q9DUB7; residues 1-720 (1 copy) + UniProt: O60684; residues 74-536 (1 copy) | | |
| AlphaFold3_Model_TTDoV_ORF1-IMPα7_0 | 0.43 | 0.51 |
| AlphaFold3_Model_TTDoV_ORF1-IMPα7_1 | 0.43 | 0.51 |
| AlphaFold3_Model_TTDoV_ORF1-IMPα7_2 | 0.43 | 0.49 |
| AlphaFold3_Model_TTDoV_ORF1-IMPα7_3 | 0.44 | 0.46 |
| AlphaFold3_Model_TTDoV_ORF1-IMPα7_4 | 0.43 | 0.18 |
| **A pTM score >0.5 means the overall predicted fold for the complex might be similar to the true structure.* | | |
| *^ipTM measures the accuracy of the predicted relative positions of the subunits within the complex; values >0.8 represent confident high-quality predictions, values <0.6 suggest likely a failed prediction.* | | |
| *Models shown in manuscript highlighted in green.* | | |

Supplementary Table S4/5. See Excel Sheet.

**Supplementary Table S6. NLSc is highly conserved in *Anelloviridae* capsid proteins, depending on protein length.** The sequences of ORF1 proteins from indicated anelloviruses were retrieved from GenBank and aligned using Clustal Omega, followed by identification of putative NLSs with cNLS Mapper. The *Anelloviridae* genera is indicated on the left, and TTDoV is indicated in orange. The C-terminal sequences are shown with the single letter amino acid code, and cNLSs are boxed. cNLSs overlapping with TTDoV ORF1 NLSc are highlighted in green, and nearby cNLSs are highlighted in purple.
